# Universal scaling relationships of marine and terrestrial photoautotrophs with differing developmental trajectories

**DOI:** 10.64898/2026.09.28.754919

**Authors:** Samuel Starko, Liam J.M. Coleman, Patrick T. Martone

## Abstract

**Backgrounds and Aims:** Partitioning of biomass to various organs is fundamental to growth and survival strategies of plant species, with broad-scale ecological implications from competition to carbon flux of ecosystems. Although partitioning patterns have been studied extensively in embryophytes, little is known about the variation and evolutionary drivers of biomass partitioning strategies in marine plants –such as seaweeds– that form marine forests analogous to those on land.

**Methods:** We measured developmental variation in standing organ biomass for 18 species of kelp and used a previously published dataset on land plants to draw comparisons between kelps and embryophytes, with respect to developmental allometry, interspecific allometry and the relationship between the two measures.

**Key results:** We report that kelps exhibit large variation in partitioning patterns, with some species differing by more than an order of magnitude in holdfast and stipe investment when controlling for size. The range of biomass partitioning patterns exhibited by kelps includes values not found among embryophytes, highlighting differences in selective pressures on land and in the ocean. However, as previously demonstrated, empirical values of interspecific scaling curves closely match those of land plants. This suggests that despite fundamentally different physical constraints, evolutionary scaling has converged on nearly identical interspecific exponents. We further demonstrate that allometric covariation exists between support organs (stipe and holdfast), highlighting the importance of physical forces in driving functional diversification of marine plants and supporting the notion that covariation of traits is a hallmark of botanical structure and function.

**Conclusions:** The results of our study illuminate the variation and evolutionary patterns of biomass partitioning in marine primary producers. Moreover, these results highlight that universal evolutionary scaling relationships can emerge from fundamentally divergent developmental and biomechanical pathways.

## Introduction

Kelps (Laminariales, Phaeophyceae) are the largest and most productive marine macroalgae in nearshore ecosystems, forming complex three-dimensional habitats that provide carbon to a diverse assemblage of coastal marine animals (Duggins et al., 1989; Steneck et al., 2002; Teagle et al., 2017). Although kelps first appeared around the Eocene-Oligocene Boundary (Kiel et al., 2024; Starko et al., 2019), they have since become the dominant foundation species on temperate reefs worldwide, diversifying into over 30 genera with distinct morphologies and varying ecological roles (Bolton, 2010). These remarkable organisms exhibit some of the fastest growth rates on Earth (Jackson, 1987), and their biomass-rich forests rank among the most productive communities globally (Pessarrodona et al., 2022). Despite often inhabiting high energy wave– or current-swept environments, kelps have specialized morphologies for resisting drag (Breitkreutz et al., 2022; Friedland & Denny, 1995; Gaylord et al., 2008; Johnson & Koehl, 1994; Koehl et al., 2008; Starko & Martone, 2016b) and can allocate biomass to support organs to prevent dislodgement (Martone, 2007; Starko et al., 2020; Starko & Martone, 2016b).

The allocation of photosynthate to various organs profoundly impacts plant ecology, influencing individual performance, community composition, and carbon cycling in ecosystems (Enquist et al., 2016). In terrestrial plants, size limitations largely arise from constraints on water transport (Niklas, 1992), with patterns of root investment linked to water availability and size-dependent geometries of the vascular system (Enquist & Niklas, 2002; Niklas & Enquist, 2001). While stem investment and branching may reflect adaptations to different environments (e.g., biotic conditions, compressive mechanical forces), maintaining water delivery necessitates specific stem and branch diameters relative to the supported leaf area (Price et al., 2007, 2009), yielding predictable biomass partitioning patterns (Niklas & Enquist, 2001, 2001; Price et al., 2007; West et al., 1999). In contrast, kelps uptake water directly through their fronds and do not transport it internally; their stipes and holdfasts function exclusively in attachment and resisting hydrodynamic forces acting on the blades. Thus, while terrestrial plant allometry is fundamentally shaped by hydraulic transport, kelp allometry arises from mechanical attachment and drag resistance, posing a powerful natural contrast in potential constraints and selective dynamics.

Flow-induced forces are believed to drive functional diversity in macroalgae, favoring a continuum of strategies that prevent mechanical failure (Martone et al., 2012; Starko et al., 2020; Starko & Martone, 2016b). Like other marine plants, kelps are mostly flexible, allowing them to reconfigure and reduce hydrodynamic loads (drag and lift) imposed by water motion (Johnson & Koehl, 1994; Martone et al., 2012; Starko & Martone, 2016b). However, unlike many other macroalgal taxa, kelps can grow in three dimensions and increase the forces they resist (Starko & Martone, 2016a; Theodorou & Charrier, 2023) through continued meristematic growth and investment in support structures (Starko & Martone, 2016a, 2016b). This combination of streamlining and mechanical reinforcement likely enables kelps to attain larger sizes than other macroalgae limited by water motion and growth axes (Martone, 2007; Wolcott, 2007). Recent findings demonstrate that interspecific scaling of organ biomass in kelps closely parallels that of terrestrial plants, suggesting that dimensional scaling may have universal impacts on the evolution of size across remarkably distantly related lineages (Cavalier-Smith, 1998; Keeling, 2004), despite the different physical phenomena shaping their morphologies (Starko & Martone, 2016a). For example, both the water requirements of embryophytes (West et al., 1997) and the drag resistance requirements of kelps relate to photosynthetic area, with water uptake scaling predictably with leaf area to ensure delivery (West et al., 1999) and flow-induced drag on kelps scaling with frond area (Starko & Martone, 2016b).

Kelp species exhibit a wide range of morphological strategies to resist wave dislodgement, with mature sizes ranging from centimeters to tens of meters (Abbott & Hollenberg, 1976). Because flow-induced forces increase with photosynthetic area, larger species may face stronger selection for investment in support structures than smaller species. Consequently, evolutionary increases in maximum size may be associated with predictable shifts in developmental biomass partitioning trajectories. In terrestrial plants, developmental biomass partitioning patterns are generally thought to mirror interspecific scaling relationships because the constraints acting on individual growth also shape evolutionary differences in size among species (Enquist & Niklas, 2002). Under this framework, interspecific scaling relationships are expected to approximate the average developmental trajectory across species. Whether this assumption also applies to kelps is unknown. Previous work has hypothesized that flow-induced forces directly limit the size of wave-swept organisms by dislodging individuals that exceed critical sizes (Carrington, 1990; Gaylord et al., 2008; Koehl, 1984; Martone & Denny, 2008; Wolcott, 2007).

Larger kelp species may therefore have evolved developmental biomass partitioning strategies that favour greater investment in support organs than smaller species, causing developmental trajectories to shift systematically with maximum size rather than remaining constant across species. If so, interspecific scaling relationships would not simply represent the average developmental strategy across species. For example, biomechanical modelling of the large kelp *Egregia menziesii* demonstrated that larger individuals have higher dislodgement risk but are stronger than necessary at all sizes (Friedland & Denny, 1995). Thus, larger species may invest in support structures early in life in anticipation of the greater hydrodynamic forces experienced later in development, increasing the likelihood of surviving to reproductive size.

Based on biomechanical theory and previous observations of drag resistance in kelps, we can pose several hypotheses about how biomass partitioning patterns may have evolved across kelps. First, we hypothesize that larger species will exhibit steeper holdfast-blade and stipe-blade developmental scaling relationships than smaller species (Fig 1a) in order to overcome size limitation effects (Carrington, 1990; Gaylord et al., 2008; Koehl, 1984; Martone & Denny, 2008; Wolcott, 2007). Second, because holdfasts and stipes function together as an integrated support system, we hypothesize that developmental investment in these organs has not evolved independently. Species that allocate more biomass to holdfasts relative to blades during growth will also allocate more biomass to stipes to ensure that attachment strength and load-bearing capacity remain mechanically matched. Third, we hypothesize that interspecific scaling relationships will differ from the average developmental trajectory of individual species if evolutionary increases in size are associated with systematic changes in biomass allocation (Fig 1a rather than Fig1b).

**Fig. 1.**
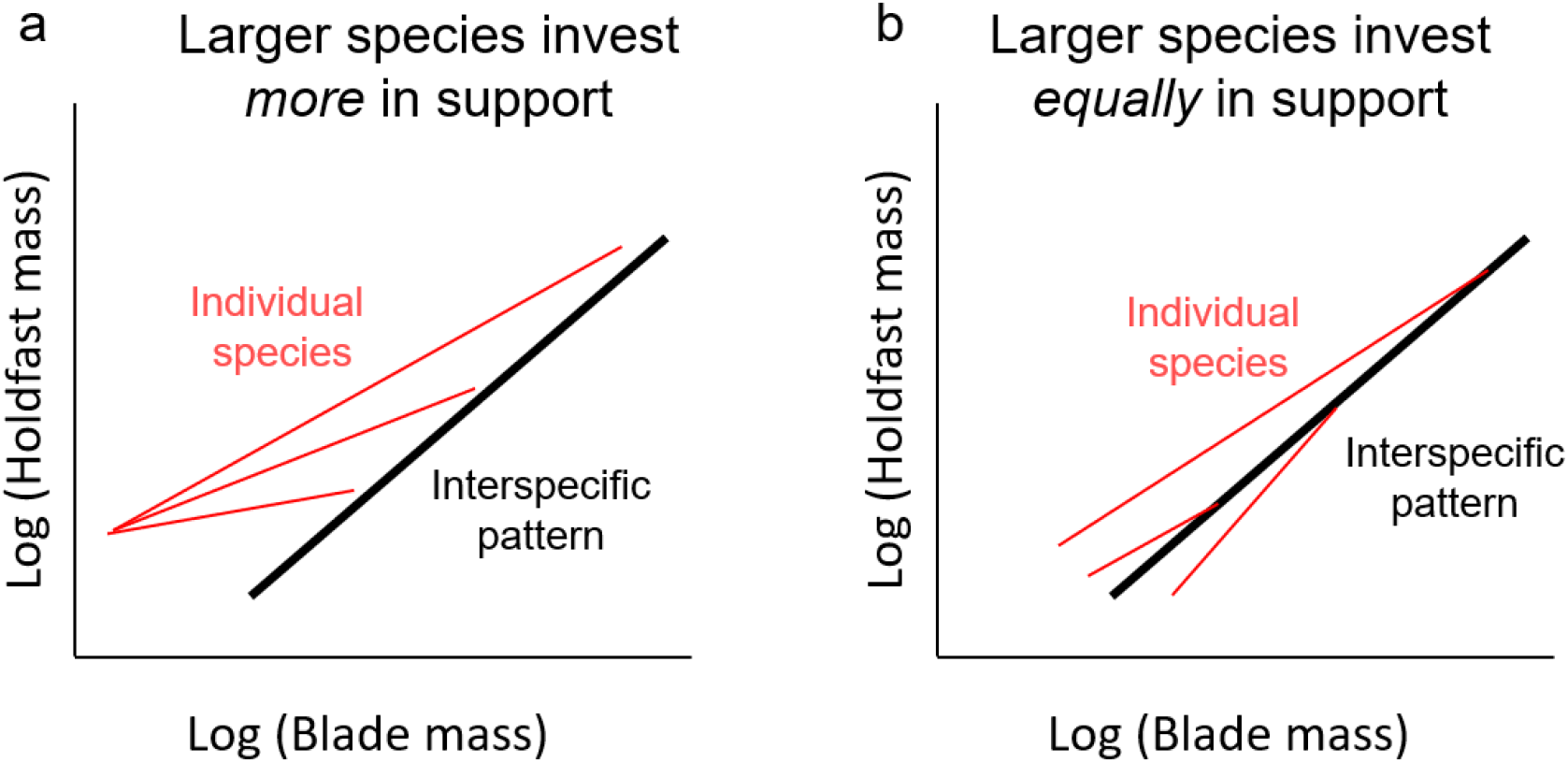
Conceptual model illustrating how developmental biomass partitioning influences evolutionary scaling relationships. Black lines represent the interspecific scaling relationship among species at maximum size, whereas red lines represent developmental trajectories of individual species. (a) If larger species progressively increase investment in support structures during development (steeper developmental scaling), the interspecific relationship does not represent the average developmental trajectory. (b) If all species exhibit similar developmental scaling regardless of maximum size, as is the paradigm for woody plants (Enquist & Niklas, 2002), the interspecific relationship approximates the average developmental trajectory.

In this study, we assess the variation in intraspecific scaling relationships of kelps by sampling several (n = 18) species from the west coast of British Columbia, a global center for kelp diversity (Starko et al., 2019). We investigate how phylogeny, morphology, and the maximum size of a species may interact to produce species-specific biomass partitioning strategies. Next, we use data from the largest individuals of each population to fit interspecific scaling curves, to investigate whether selective pressures influence the evolution of size and biomass partitioning strategies. Finally, we compare the patterns observed across the kelps to those found in land plants by fitting both intra– and interspecific scaling relationships to a large dataset of embryophytes (from Poorter et al., 2015). We specifically aim to address four questions: (1) What range of biomass partitioning strategies have kelps evolved to adapt to diverse environments? (2) How do phylogenetic relationships influence developmental biomass allocation patterns among species? (3) Are evolutionary increases in maximum size associated with predictable shifts in developmental allometric trajectories? and (4) How do developmental and interspecific scaling relationships of kelps compare with those of embryophytes?

## Materials and Methods

### Allometric approach

Biomass partitioning patterns between organs were modeled as allometric power scaling relationships, such that:

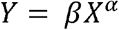

where Y and X are the masses of two organs, β is the allometric or scaling constant (absolute value or intercept of the relationship), and α is the allometric or scaling exponent. In order to linearize these relationships, data were log transformed, such that:

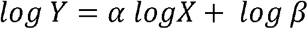

Thus, α, the scaling exponent is equal to the slope of the log-log linear relationship between organ masses.

### Sample Collection

Samples of 18 kelp species were collected for allometric analyses from various sites along the west coast of British Columbia (Bamfield, Calvert Island, Vancouver, and Victoria; Table 1). An effort was made to collect samples that spanned the entire range of sizes at each site. Samples from all sites were separated into blade(s), stipe(s), and holdfast and each organ was weighed separately. Kelps from Barkley Sound, Vancouver and Calvert Island were air-dried at room temperature for at least 12 hours prior to drying at higher temperatures. All but the largest of these kelps were placed in a drying oven at 60 ° C for at least 24 hours. To determine appropriate drying time, subsamples were weighed at intervals until dry weight had stabilized. The largest *Egregia* and *Macrocystis* samples were dried at 37-39 ° C for 24 hours in an industrial kelp drier (Canadian Kelp Resources, Bamfield, BC). Large kelps collected from Victoria were sun-dried for 24 hours (over the course of two days) and then dried in a small room that was heated by four space heaters. The room was kept at 32 ° C for 48 hours and then 37-39 ° C for 18 hours. To determine whether the methods used to dry the largest kelps were effective compared to traditional 60 ° C drying oven techniques, subsamples were placed in a 60 ° C oven for 24 hours following each of these protocols. Error generated from different drying methods was generally 2-3% (always less than 4.5%), which was considered inconsequential given the large range of biomasses and the fact that data were log-transformed for purposes of analysis.

**Table 1.** Collection information for species sampled in this study.

| Species | Morphology | Sample Size | Habitat | Site |
| --- | --- | --- | --- | --- |
| *Lessoniopsis littoralis* | Stipitate | N = 36 | Intertidal | Brady's Blowhole, Bamfield |
| *Hedophyllum sessile* | N/A* | N = 37 | Intertidal | Brady's Blowhole, Bamfield |
| *Postelsia palmaeformis* | Stipitate | N = 24 | Intertidal | Cape Beale, Bamfield |
| *Laminaria setchellii* | Stipitate | N = 36 | Intertidal | Eagle Bay, Bamfield |
| *Ecklonia arborea* | Stipitate | N = 20 | Intertidal | Eagle Bay, Bamfield |
| *Egregia menziesii* | Canopy | N = 21 | Intertidal | Eagle Bay, Bamfield |
| *Macrocystis pyrifera* | Canopy | N = 19 | Intertidal | Eagle Bay, Bamfield |
| *Alaria marginata* | Prostrate | N = 31 | Intertidal | Eagle Bay, Bamfield |
| *Laminaria ephemera* | Prostrate | N = 14 | Intertidal | Edward King Island, Bamfield |
| *Saccharina latissima* | Prostrate | N = 24 | Intertidal | Kitsilano Beach, Vancouver |
| *Laminaria yezoensis* | Prostrate | N = 19 | Intertidal | West Beach, Calvert Island |
| *Agarum fimbriatum* | Prostrate | N = 20 | Subtidal | Bamfield Inlet, Bamfield |
| *Costaria costata* | Prostrate | N = 29 | Subtidal | Ogden Point, Victoria |
| *Nereocystis luetkeana* | Canopy | N = 26 | Subtidal | Ogden Point, Victoria |
| *Pleurophycus gardneri* | Stipitate | N = 29 | Subtidal | Ogden Point, Victoria |
| *Pterygophora californica* | Stipitate | N = 22 | Subtidal | Ogden Point, Victoria |
| *Hedophyllum nigripes* | Prostrate | N = 21 | Subtidal | Ogden Point, Victoria |
| *Cymathaere triplicata* | Prostrate | N = 21 | Subtidal | Ogden Point, Victoria |

\*Indicates the unusual morphology of *H. sessile* which lacks a stipe but is otherwise prostrate.

### Data on embryophytes

Data on embryophyte biomass partitioning were taken from a comprehensive review that included a database of organ biomass observations (Poorter et al., 2015), including those of the well-known Cannell (1982) dataset. We omitted species from the analysis that had few replicates (N < 15) and/or spanned less than two orders of magnitude in leaf biomass. *Cunninghamia lanceolata* was also omitted from the analysis because data for these species had strong heterogeneity of variance. The resulting dataset contained developmental organ partitioning data that was appropriate for fitting allometric relationships for 113 embryophyte species filtered from the initial dataset of ∼11,000 measurements.

### Statistical analysis

All statistical analyses and visualizations were performed in R (R Core Team., 2024) (https://www.r-project.org/). All regression analyses involving organ partitioning were performed using reduced major axis (RMA) slopes of log-log data using the “lmodel2” package (Legendre, 2014). This statistical technique is often used in allometric analyses since it aims to minimize residual size across both axes (see Niklas 1994). Because allometric constants are arbitrarily defined as the intercept of log-log regressions, their values depend largely on the unit of measure. Here, data were analyzed and presented using centigrams, since 1 cg is the approximate size of the smallest individuals that were collected from most kelp species. Thus, intercepts in this study represent an initial value of organ investment rather than an arbitrary point along the regression.

We analyzed data for each species separately to identify developmental organ biomass scaling relationships and then tested for differences between species by comparing 95% confidence intervals of exponents and constants. To identify interspecific scaling relationships associated with evolutionary differences in size, we calculated the average organ biomasses from the three largest individuals of each species. Although this may not be the absolute maximum size of a species, this was intended to be representative of the largest individuals at the sites sampled. Thus, this represents a local ecological maximum rather than an absolute species maximum, providing a consistent and conservative basis for interspecific comparison. Using these species-level averages, we then fit allometric models (as above) to estimate interspecific scaling relationships. We also tested for differences between interspecific exponents and constants in kelps and embryophytes using the “smatr” package in R (Warton et al., 2012).

Morphologically, species were binned into prostrate, stipitate, and canopy categories based on their position within kelp communities and stature of their stipe(s). *Hedophyllum sessile* was not assigned to one of these categories, since it lacks a stipe at maturity. We did not use these bins for any statistical analysis, only for visualization.

We used a previously published time-calibrated phylogeny (Starko et al., 2019) for phylogenetic comparative methods involving allometric parameters. We included *Laminaria yezoensis* into the tree by editing the tree file manually to include a topology inferred from a past study on the genus *Laminaria* (Rothman et al. 2017). We tested whether there was a phylogenetic signal on any of the scaling parameters by calculating Blomberg’s K and Pagel’s Lambda using the “phylosig” package in R.

## Results

Species differed significantly in exponents and intercepts of both holdfast and stipe partitioning allometries (Fig. 2) with exponents ranging between 0.56 and 1.29 for both holdfast and stipe biomass relative to blades. Seasonally ephemeral species *Laminaria ephemera* and *Cymathaere triplicata* had the lowest rates of holdfast investment with exponents of 0.56-0.57, meaning that holdfasts only accumulated biomass at a rate approximately square root of that of the blades. In contrast, many of the largest (e.g., *Egregia menziesii, Macrocystis pyrifera*) and most arborescent species (e.g., *Postelsia palmaeformis, Pterygophora californica*) had the highest rates of holdfast allocation with exponents equal to unity (i.e., linear relationship between holdfast and blade biomass) or well above (e.g., = 1.29) and with intercepts that were also high relative to other species. *Hedophyllum sessile*, which had the lowest holdfast-blade scaling constant also had the greatest holdfast-blade exponent suggesting that at small sizes this species has a particularly small holdfast relative to its blade but that it has the greatest rate of holdfast biomass accumulation of any kelp examined here.

**Fig 2.**
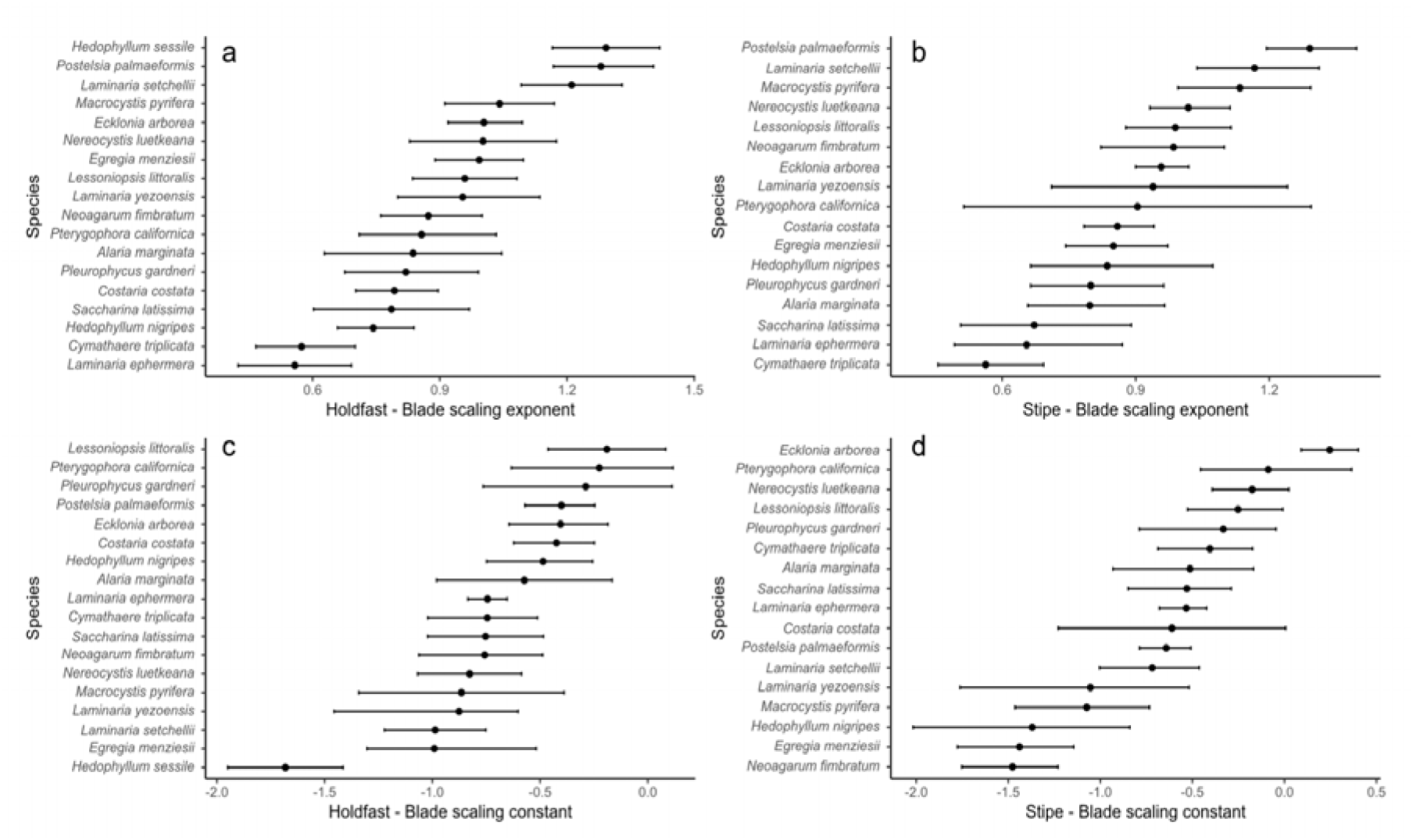
Organ biomass scaling relationships of kelp species. (a,c) Holdfast-blade relationships; (b, d) Stipe-blade relationships. Bars represent 95% confidence intervals of estimated slope and intercept of allometric models. Note that stipe relationships exclude *H. sessile* which lacks a stipe. All models are RMA regressions.

There was no significant effect of phylogeny on either holdfast-blade (Lambda < 0.01; P = 0.99; Lambda = 0.60; P = 0.68) or stipe-blade scaling exponents (Lambda < 0.01; P = 0.99; K = 0.67; P = 0.23), indicating high evolutionary lability in these measures (Fig. 3). However, there was a significant phylogenetic signal for stipe-blade scaling constants (Lambda = 0.93; P = 0.008; K = 1.14; P = 0.003) and holdfast-blade scaling constants (Lambda = 0.69; P = 0.09; K = 0.97; P = 0.02), suggesting that there may be evolutionary constraints on biomass allocation early in life (Fig. 3). Several closely related species differed substantially in both scaling exponents and scaling constants (Fig. 4). For example, members of the giant kelp clade (*Macrocystis, Nereocystis* and *Postelsia*) exhibited markedly different developmental trajectories despite their close evolutionary relationships. Similarly, three species of *Laminaria* differed considerably in both holdfast and stipe allocation patterns, with the ephemeral species *Laminaria ephemera* exhibiting substantially lower support investment than the perennial species *L. setchellii* and *L. yezoensis*. However, three closely related species from the Alariaceae, *Pterygophora, Pleurophycus*, and *Lessoniopsis*, exhibited relatively similar allometric intercepts (Fig. 3), indicating weak but localized phylogenetic signal in developmental biomass allocation.

**Fig 3.**
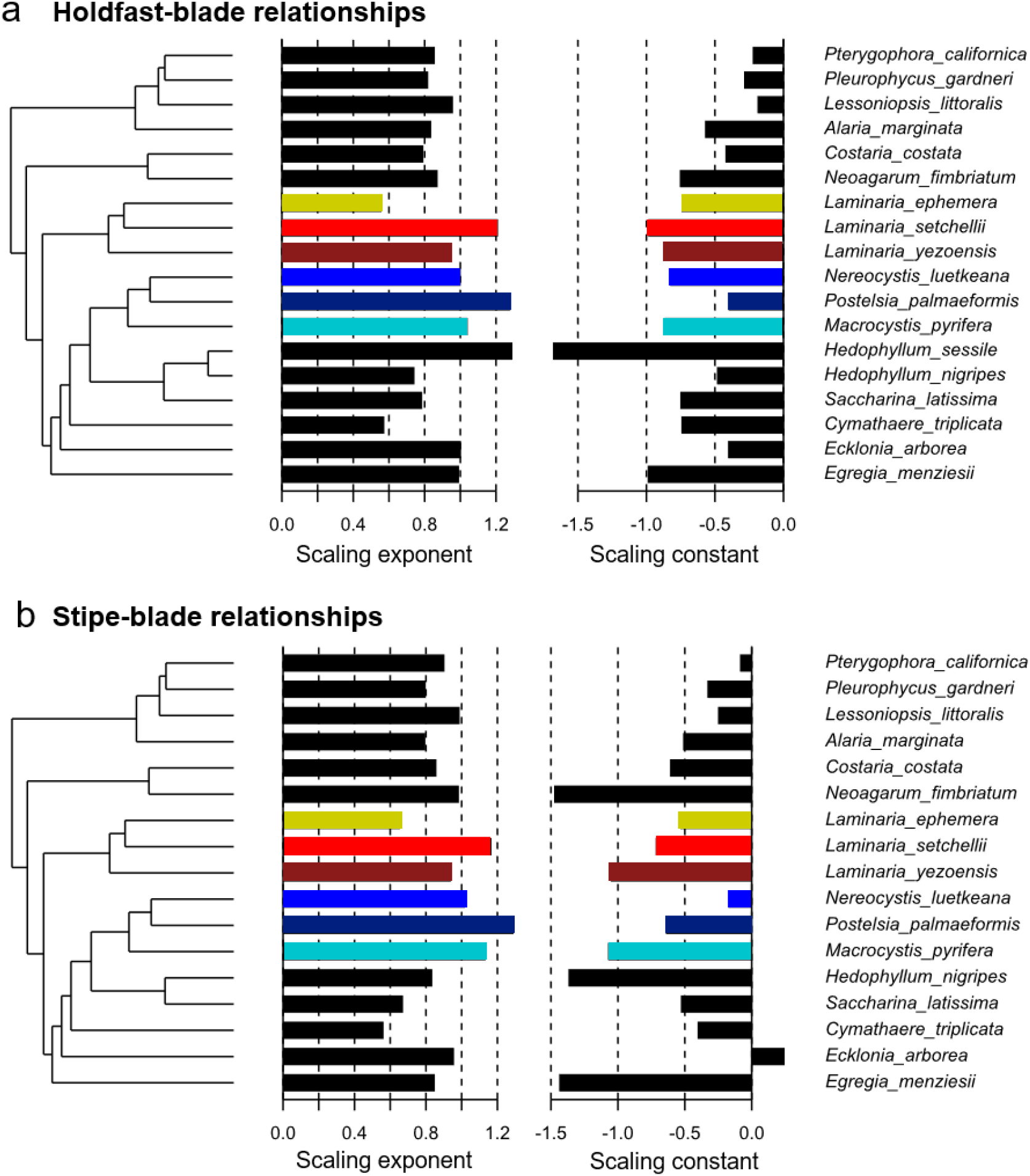
Phylogenetic relationships between kelp species and their allometric parameters. (a) Holdfast-blade relationships; (b) Stipe-blade relationships. Both exponents and constants are scaled and centred for visualization. Note that stipe relationships exclude *H. sessile* which lacks a stipe. Colours indicate the focal species groups highlighted in Fig. 4 to facilitate comparison between phylogenetic position and developmental allometric relationships.

**Fig 4.**
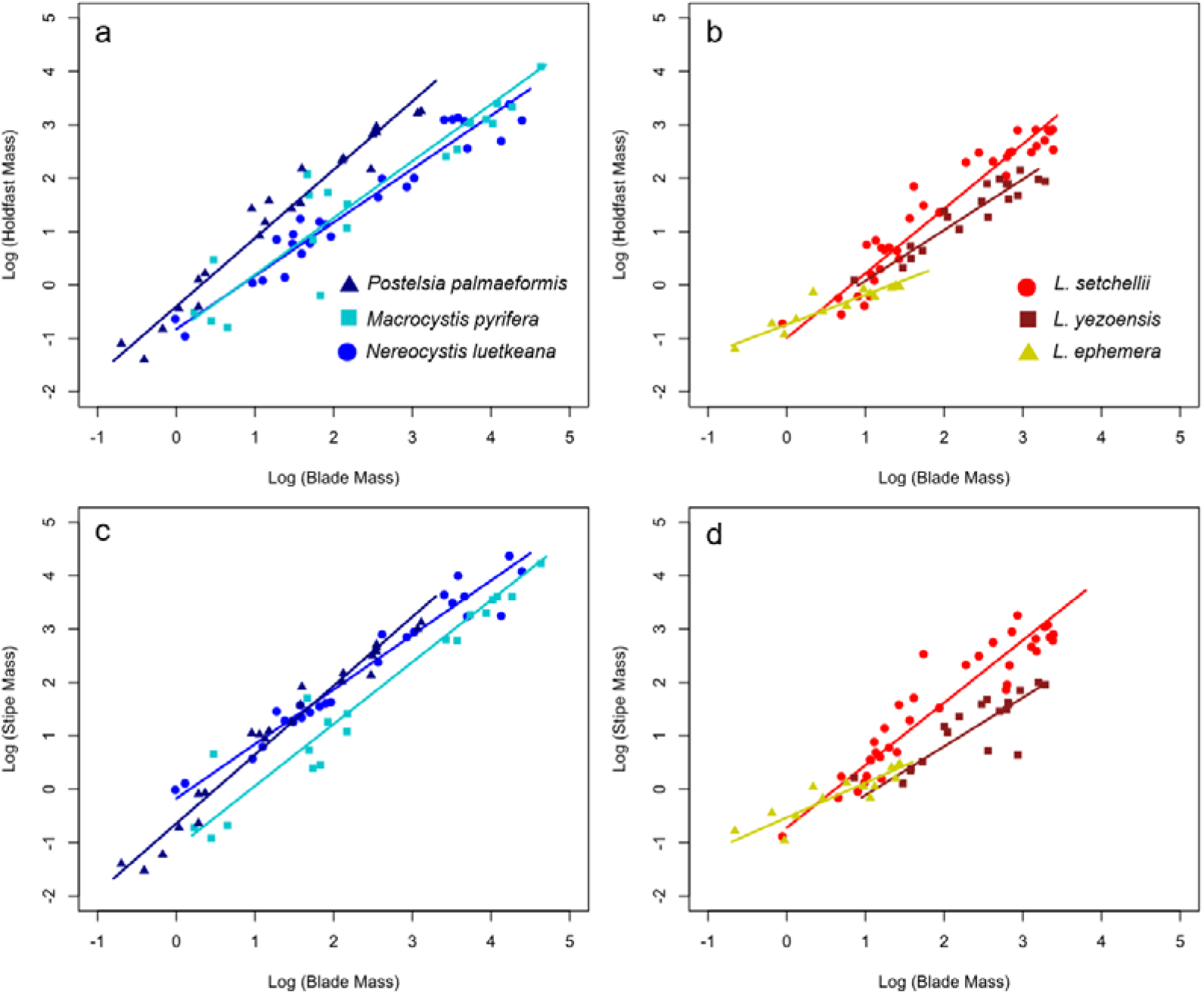
Examples of developmental allometric relationships between organs for groups of closely related species with different habitats or life-histories. (a,c) The giant kelp clade including *Postelsia palmaeformis, Macrocystis pyrifera* and *Nereocystis luetkeana.* (b, d) Three species of *Laminaria: L. setchellii, L. yezoensis, L. ephemera*.

Stipe and holdfast scaling exponents and constants covaried across species (Fig 5; Linear model [exponents]: t = 9.445, p < 0.001, Linear model [constants]: t = 2.384, p < 0.05) and morphology played an important role in determining where species fell along the covariance continuum. In particular, stipitate and canopy kelps tended to have higher stipe-blade and holdfast-blade scaling exponents than prostrate kelps (Fig 5). In contrast, scaling constants only covaried weakly, with large variation between species, indicating that the overall magnitude of investment may differ somewhat between stipe and holdfast but that these organs scale precisely as they get larger. Both holdfast and stipe scaling exponents significantly varied positively with size (Linear model [holdfast]: t = 2.124, P < 0.05, Linear model [stipe]: t = 2.139, P < 0.05).

**Fig 5.**
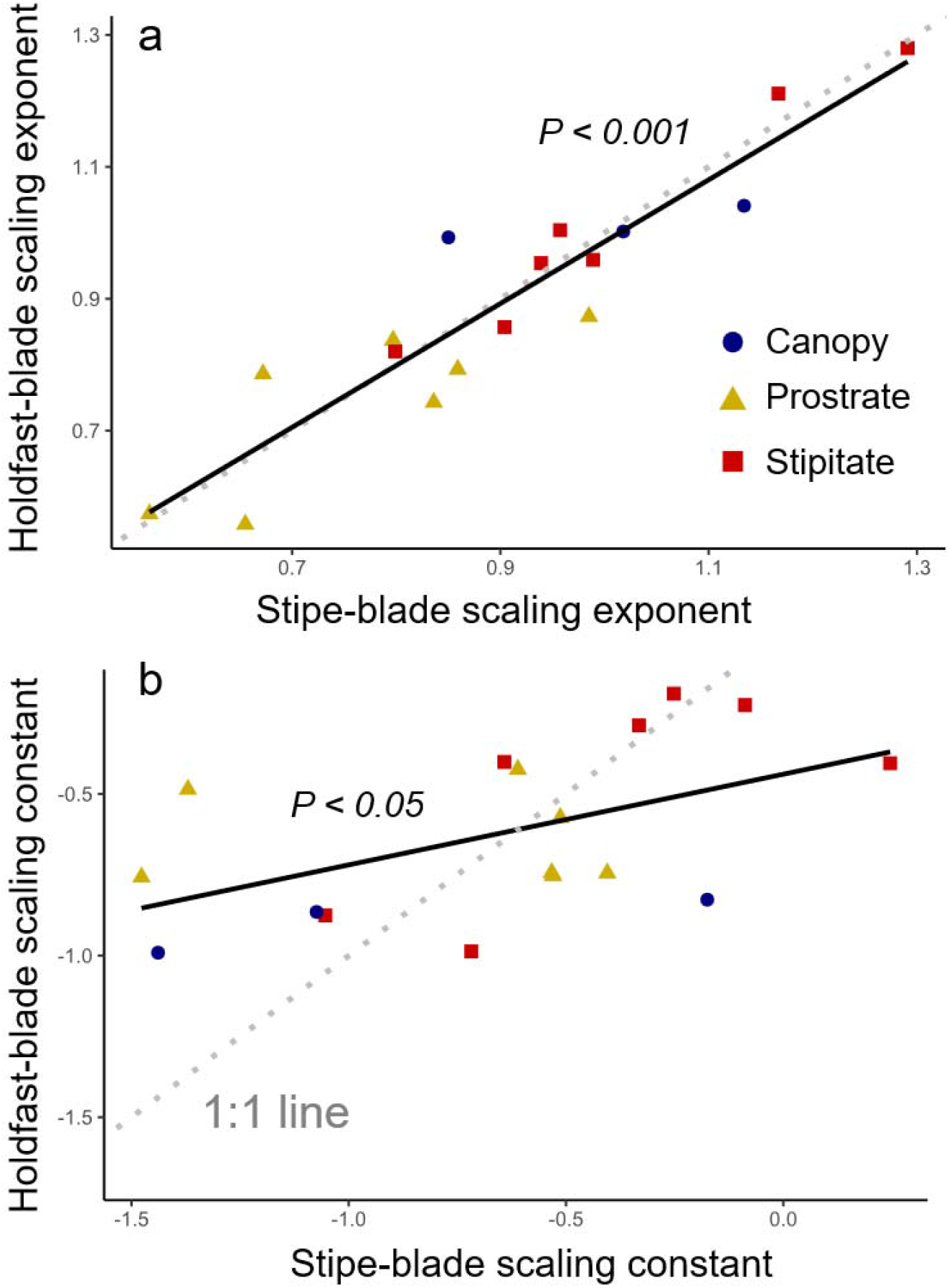
Covariation in scaling relationships between holdfast and stipe allocation. (a) Strong positive relationship between scaling exponents. (b) Weak positive relationship between scaling constants. Dotted lines indicate 1:1 while solid lines indicate best fit regression lines. Note that plots exclude *H. sessile* which lacks a stipe.

Comparing scaling relationships of kelps to those of embryophytes, we found that there was extensive overlap but also key differences in developmental patterns. Although scaling exponents largely overlapped across the two groups, there were also values unique to both kelps and embryophytes. While embryophyte root-leaf scaling exponents varied from 0.9 to 1.46, equivalent scaling exponents across the kelps range from 0.56 to 1.29 indicating lower minimum values in kelps, as well as a slightly larger range of values (Fig 6). To compare developmental and evolutionary scaling patterns, we contrasted the distribution of developmental (intraspecific) scaling parameters with the parameter values estimated from interspecific scaling relationships fit using the largest individuals of each species (dotted lines in Fig 6).

**Fig 6.**
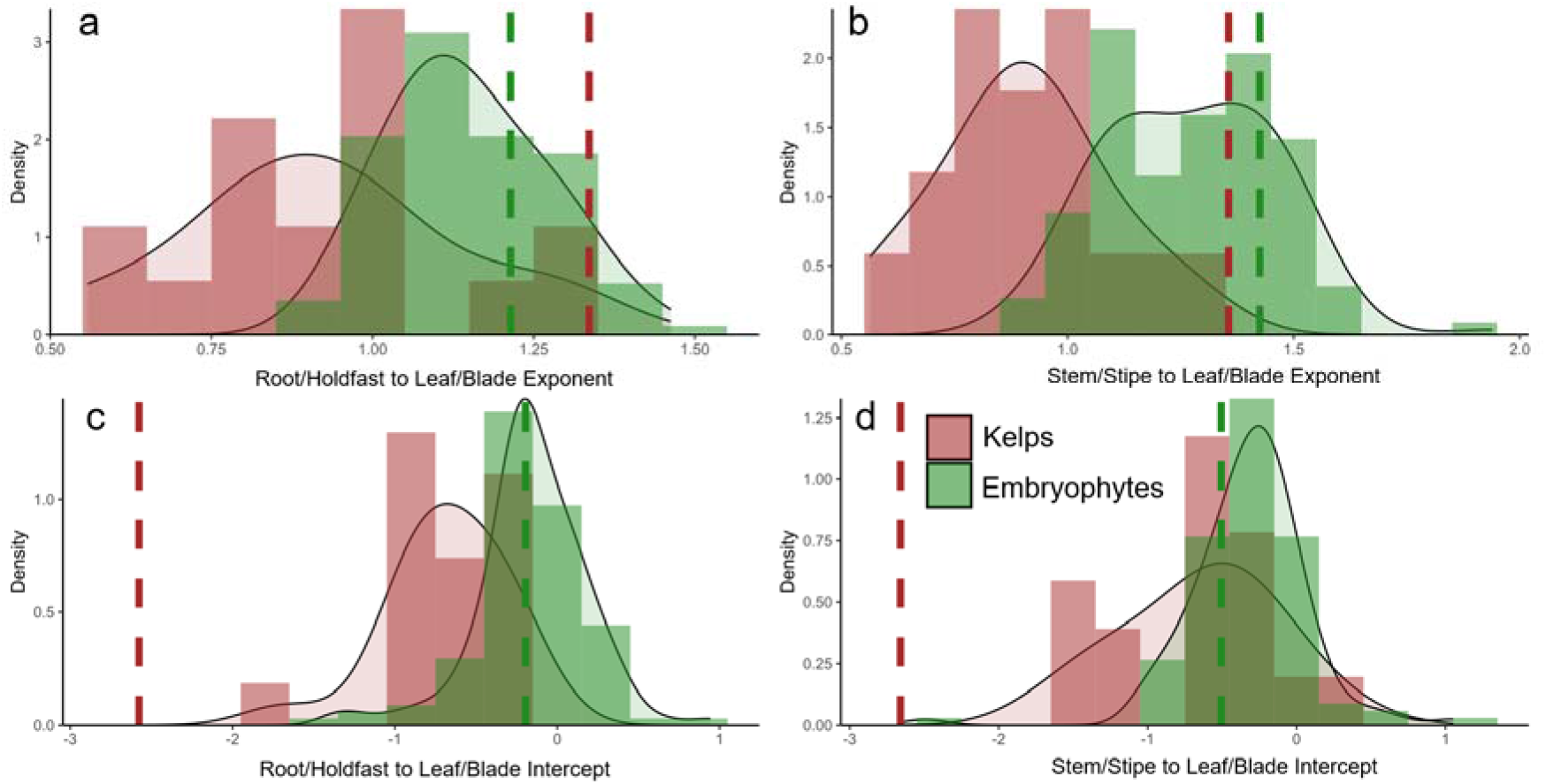
Distribution of developmental (intraspecific) scaling relationships for kelps (n = 18 species) and land plants (n = 113 species). Dotted lines indicate the exponent (a,b) or constant (c,d) estimated from interspecific scaling relationships fit using the largest individuals of each species (Figs. 8,9). Not that stem relationships exclude *H. sessile* which lacks a stipe.

While there was a positive relationship between maximum size and developmental scaling exponent for both groups, this relationship was much steeper for kelps, particularly for holdast-blade/root-leaf relationships (Fig. 7), reflecting a sharp increase in holdfast and stipe investment with increasing size. Because developmental scaling exponents varied systematically with maximum size, the interspecific exponent estimated from the largest individuals of each species did not correspond to the mean developmental exponent across species (Figs 6, 8). Instead, the interspecific exponent (dotted lines in Fig. 6) was offset from the centre of the developmental exponent distributions in both groups, but to a much greater degree in kelps. This discrepancy reflects the stronger association between size and developmental allometry in kelps than in embryophytes (Fig. 7). Despite these differences, scaling exponents were nearly identical across embryophytes and kelps when considering interspecific relationships among the largest individuals of each species (Fig. 9). For both root-leaf and stem-leaf relationships, there were no significant differences in exponent between kelps and embryophytes (SMA slope tests: P > 0.05). However, intercepts were significantly lower for kelps (SMA intercept test [root-leaf]: P < 0.001; SMA intercept test [stem-leaf]: P < 0.001), indicating a lower overall proportion of biomass in stipes and holdfasts of kelps than in stems and roots of embryophytes, despite similar interspecific scaling exponents.

**Fig 7.**
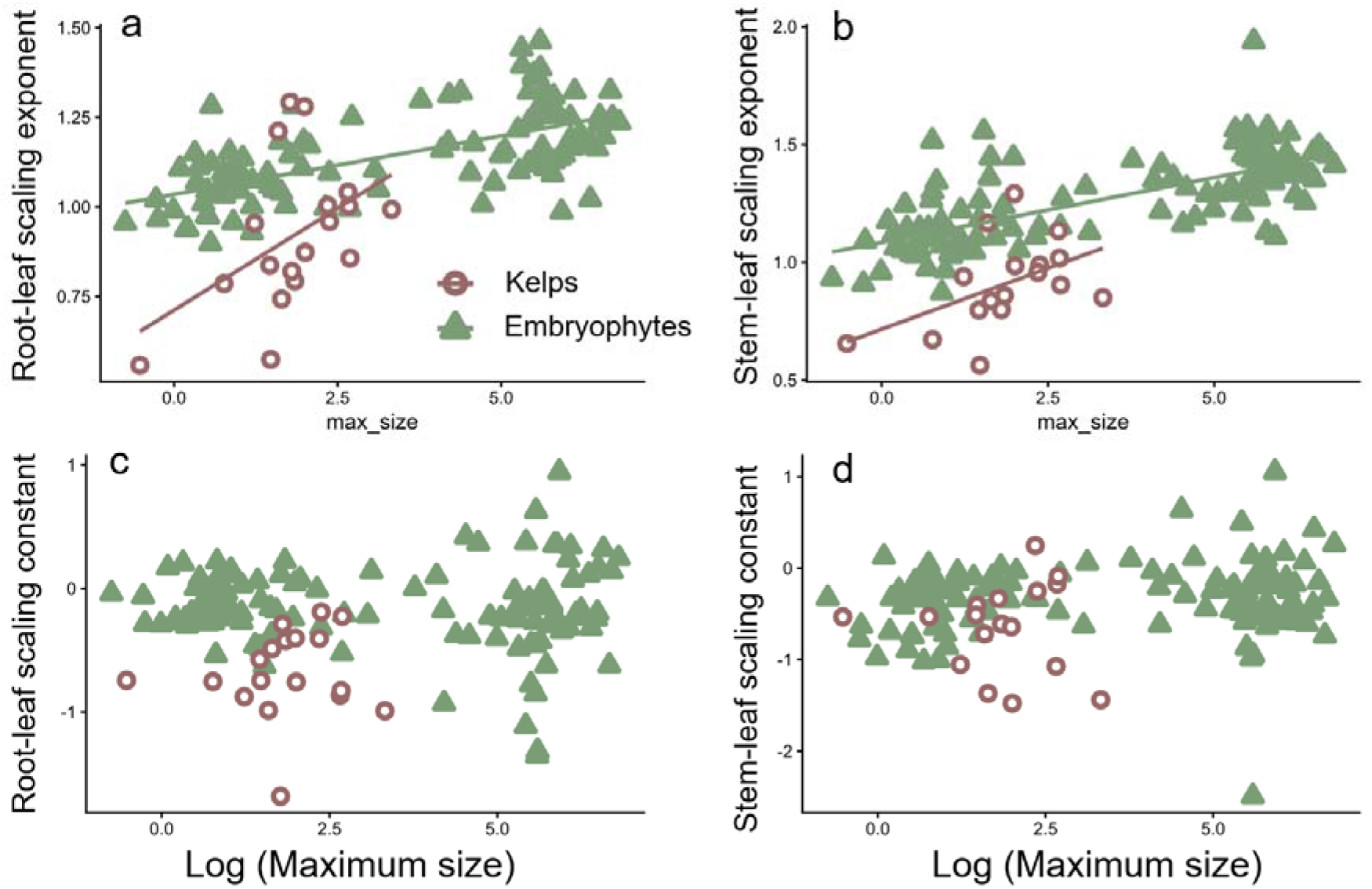
The relationship between size and allometric scaling relationships for both embryophytes and kelps. Regression lines indicate significant relationships (P < 0.05). Note that root-leaf and stem-leaf relationships for embryophytes are equivalent to holdfast-blade and stem-blade relationships in kelps, respectively.

## Discussion

Our study provides the first quantitative analysis of developmental organ biomass scaling across a diversity of kelp species. The results reveal remarkable variation in how kelps partition resources to their support organs (holdfasts and stipes) relative to their blades. Scaling exponents ranged from ∼0.5 to ∼1.3 for both holdfast and stipe investment, indicating some species invest minimally in support relative to photosynthetic area, while others allocate disproportionately more across support organs as they grow larger. This variability in biomass partitioning in kelps is marginally greater than that observed across terrestrial embryophytes (0.9-1.46), despite far fewer species in the kelp dataset (18 kelp species vs 113 embryophyte species).

Our first prediction was that larger species would exhibit steeper holdfast– and stipe-blade scaling relationships because larger individuals experience disproportionately greater hydrodynamic loading. Within kelps, biomass partitioning strategies were highly variable, reflecting differences in morphology of different species. However, the largest kelp species did indeed invest more in holdfasts and stipes through development than small species (e.g., *Laminaria ephemera*), likely as a prerequisite to survive wave forces experienced at large sizes (Friedland & Denny, 1995; Gaylord et al., 2008; Starko & Martone, 2016b). This manifested as a significant relationship between holdfast– and stipe-blade scaling relationships and the maximum size of species. Larger species are also often perennial (with clear exceptions; e.g., *Nereocystis luetkeana*) and some of which may not reproduce in their first year (Bartsch et al., 2008; L. Druehl pers comm). Thus, longer-lived species might invest more in support as a means of increasing mechanical safety and ensuring survival to reproductive age. Moreover, stipitate and canopy-forming morphologies exhibited higher scaling exponents than flexible, prostrate species (Fig 5) that can likely reconfigure more easily and reduce the flow-induced forces they experience (Starko & Martone, 2016b).

Interestingly, both kelp and embryophyte scaling exponents increased with increasing maximum size. This may reflect similarities in selective pressures associated with evolving overall greater biomasses. While some diminutive kelps invest very little in holdfasts and stipes (akin to ephemeral terrestrial herbs that invest little in roots and stems), larger canopy-forming and arborescent species partition substantially more biomass in holdfasts to withstand wave forces (Friedland & Denny, 1995; Johnson & Koehl, 1994; Starko & Martone, 2016b), analogous to trees investing disproportionately in trunks and roots as they grow larger (Enquist et al., 2007; Poorter et al., 2015).

Compared to embryophytes, the impact of size on developmental scaling was much steeper in kelps, however. While larger terrestrial species tend to have greater stem or root scaling relationships (Poorter et al., 2015; Fig 7), kelps appear to have undergone more dramatic developmental shifts as they have evolved increasing size, drastically increasing support expenditure to compensate for heightened drag. For kelps, the largest species may exhibit elevated early relative investment in support structures, consistent with anticipatory mechanical reinforcement. This frontloading of investment early in ontogeny has been previously observed for the large kelp, *Egregia menziesii* (Friedland & Denny 1995).

Phylogeny did not significantly predict biomass partitioning exponents, suggesting high evolutionary lability in rates of biomass allocation. This is consistent with past work demonstrating high lability in morphological and mechanical traits with minimal phylogenetic signal (Starko et al., 2020). Even closely related species could differ markedly, though some phylogenetic signal was apparent across the Alariaceae and there was a weak phylogenetic signal on scaling constants, suggesting that early life stage allocation may be under some level of evolutionary constraint. Consequently, while there might be some evolutionary constraint on biomass partitioning, rates of support investment are highly labile and likely play an important role in the adaptive evolution of taxa within the kelps.

Our second prediction was that support organs (holdfast and stipe) would evolve in concert rather than independently. Indeed, we found strong covariation between stipe and holdfast scaling exponents regardless of phylogeny, providing compelling evidence that an integrated response to wave-induced forces drives the scaling of supportive organs in kelps. Species that allocate proportionally more biomass to holdfasts through development also allocate proportionally more biomass to stipes, suggesting that attachment strength and load-bearing capacity evolve as a coordinated biomechanical system. Notably, this relationship was much stronger for scaling exponents than for scaling constants (although both relationships were significant). This difference is consistent with the expectation that developmental trajectories of support investment should remain mechanically matched, whereas the absolute magnitude of investment may be influenced by morphological features such as stipe length, blade architecture, or the presence of buoyant floats derived from stipe tissue. Thus, species can differ substantially in the overall proportion of biomass allocated to support structures while maintaining similar developmental scaling relationships between the two organs. More broadly, this finding is consistent with the idea that trait covariation is fundamental to plant form and function (Price et al., 2007).

Our final prediction was that interspecific scaling relationships would not correspond to the average developmental trajectory if evolutionary increases in size were accompanied by shifts in developmental allocation. Despite pronounced differences in developmental scaling across kelps and embryophytes, the resultant organ scaling relationships when examining full-grown individuals of each species (i.e., interspecific scaling relationships) are remarkably similar (as observed in Starko & Martone 2016). If interspecific scaling relationships simply represented the average developmental strategy across species, the corresponding parameter estimates would be expected to fall near the center of the developmental distributions shown in Fig. 6. Instead, interspecific exponents and constants were consistently offset from the mean developmental values, particularly in kelps, indicating that evolutionary increases in size are associated with systematic shifts in developmental trajectories (see above). For kelps, the vast majority of datapoints from smaller-than-maximum individuals fell above the interspecific holdfast-blade and stipe-blade relationships (Fig 8). This is reflected in the difference between intra– and interspecific scaling exponents and constants in Fig 6. Thus, interspecific relationships in kelps likely represent a minimum holdfast investment for a given size that can avoid dislodgement with developmental trajectories all falling on the “safe” side of this curve. For embryophytes, interspecific relationships are much closer to the mean of developmental strategies but do imply greater investment in roots than expected by this average, similar to kelps. In both cases, interspecific relationships represent the outcome of selection for increased size, where smaller species may avoid unnecessary investment in support structures while larger species require investment to avoid mechanical or hydraulic failure. For land plants, previous work using this same dataset found that the exponent of organ scaling relationships changes when moving from small to large datapoints (Poorter et al., 2015). Here, we explicitly separated out interspecific patterns from developmental trajectories and found that interspecific relationships between embryophyte species are actually log-log linear across >7 orders of magnitude in plant biomass, whereas developmental organ scaling exponents change significantly with increasing size.

**Fig 8.**
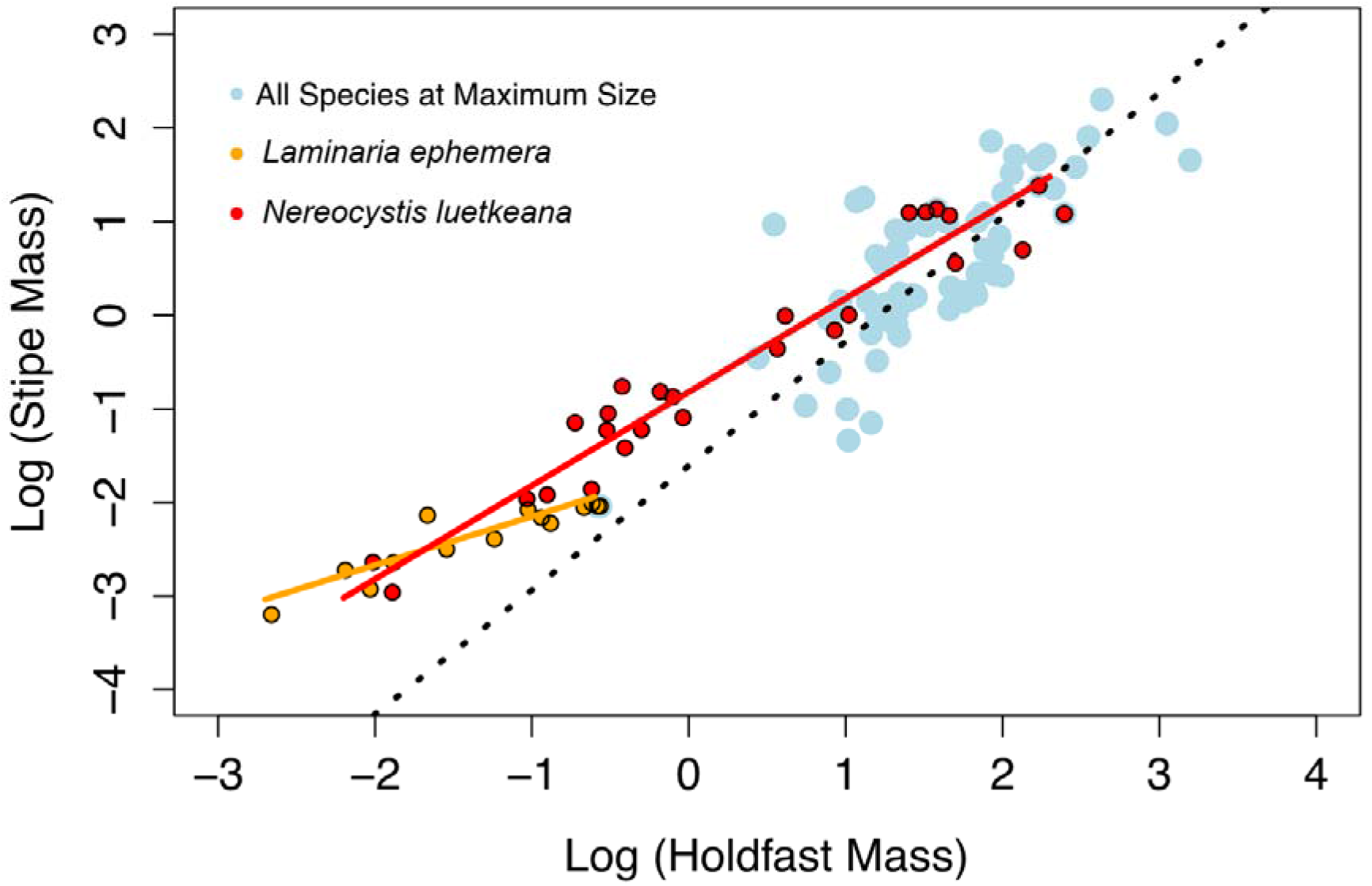
Examples of how developmental scaling relationships relate to interspecific scaling relationships for kelps. Note that developmental curves fall above the interspecific relationship.

**Fig 9.**
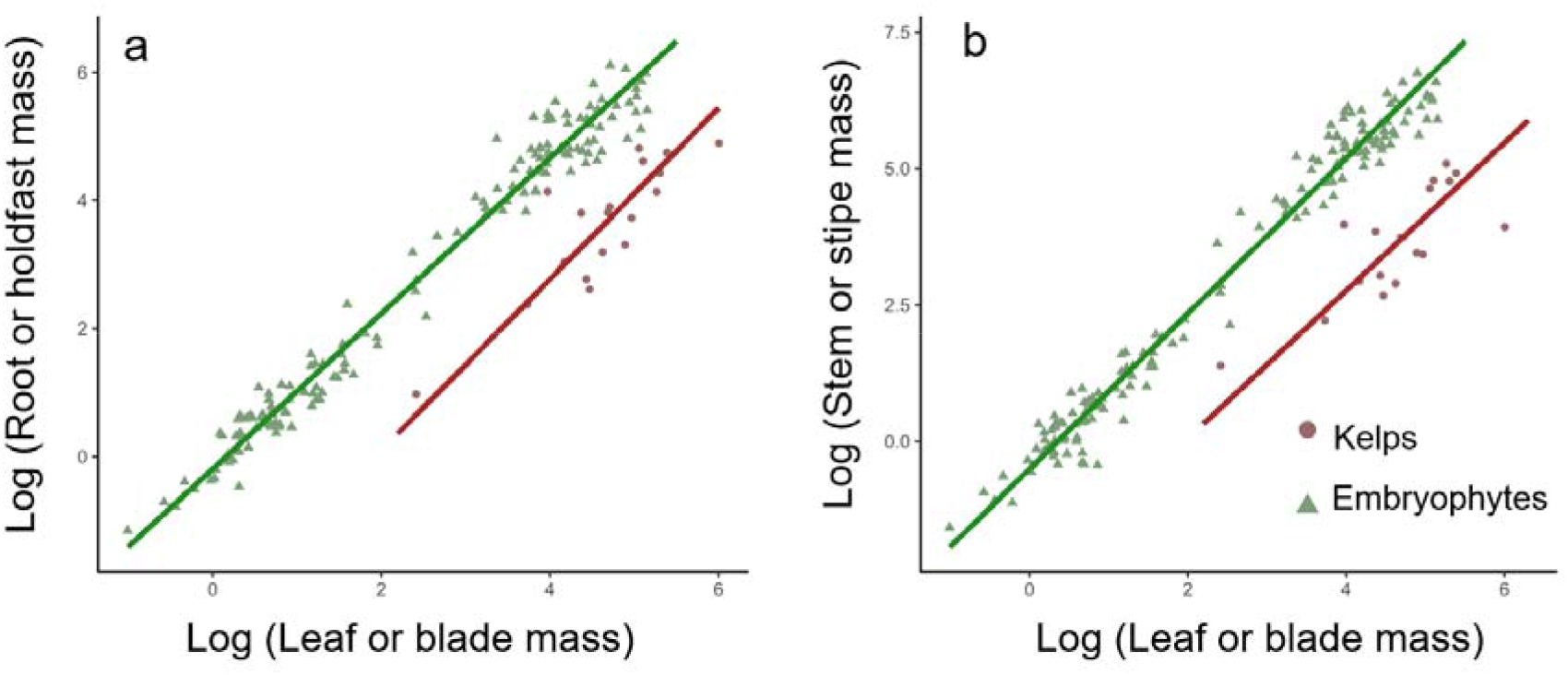
Interspecific scaling relationships between full-grown individuals (i.e., individuals at maximum size; n = 3) of each species for kelps and embryophytes. Lines represent RMA regressions fit to each dataset. In both plots, slopes were not significantly different (SMAtest: p < 0.05) but intercepts were (SMAtest: p < 0.001).

While this work provides insight into evolutionary patterns of biomass allocation in marine plants, many intriguing questions remain. For instance, how do intraspecific developmental patterns of a species vary across wave exposures? To what extent are developmental scaling relationships determined genetically versus by plastic responses to the environment? How does holdfast investment impact overall productivity, given metabolic costs (Arnold & Manley, 1985)? More explicit integration of size-dependent survival, reproduction and biomechanical limits could help elucidate the ecological drivers of observed partitioning strategies. Expanded field sampling to other parts of species’ ranges or to other parts of the world could help to capture more variation within the kelp lineage and identify geographical differences in selective pressures. Indeed, genomic work has also revealed distinct lineages within some of the species sampled here (e.g., the *Alaria marginata* complex; (Bringloe et al., 2021)), underscoring the value of confirming lineage identity when extending allometric comparisons across its range. Moreover, experimental approaches that manipulate flow and quantify organ investment could help shed light on these remaining uncertainties. Nonetheless, this study establishes the range of strategies used by one of the most prolific clades of marine habitat formers to navigate the universal challenge of allocating limited resources across integrated functions.

### Conclusions

Overall, this study highlights universal constraints on optimal biomass partitioning shared across phototrophs and also key differences in kelps and embryophytes, which are distantly related and have evolved under divergent selective regimes (Niklas & Enquist 2001, Enquist & Niklas 2002, Price et al. 2007). Substantial variation in partitioning strategies has likely contributed to the functional diversification of kelps, allowing them to occupy a wide variety of coastal habitats worldwide. Our investigation reveals remarkable diversity in the biomass partitioning strategies employed by kelp species. While following similar overall interspecific patterns as terrestrial plants, kelps exhibit a broader range of scaling exponents, likely reflecting differences in selective pressures on land versus in the ocean. Substantial work remains to fully understand the biomechanical, ecological, and evolutionary drivers leading to the observed spectrum of partitioning strategies and their subsequent impact on individual performance and community dynamics.

## Acknowledgements

The authors acknowledge L. Liggan, R. Munger, and B. Radziej for help in the field and laboratory and to L. Druehl and Canadian Kelp Resources for allowing use of their industrial kelp drier. Thanks to E. Clelland, S. Gray and the rest of the BMSC staff for support. Funding for this project was provided by the Natural Sciences and Engineering Research Council (NSERC) Discovery Grant to P.T.M. as well as a CGSM and PGSD to S.S. S.S. was also funded by Killiam Trusts and the Forrest Research Foundation. Additional travel funding and accommodation was provided by the TULA Foundation.

